# 3D co-cultures of primary human hepatocytes and Kupffer-like cells to address innate immune responses to rAAV

**DOI:** 10.1101/2025.07.08.663523

**Authors:** Isabella Ramella Gal, Francisca Arez, Inês P. Correia, Giacomo Domenici, Sofia Fernandes, Gabriela Silva, Inês Saldanha, Nadia Duarte, Catarina Freitas, Paula M. Alves, Udo Maier, Ana Sofia Coroadinha, François Du Plessis, Catarina Brito

## Abstract

Recombinant adeno-associated viruses (rAAVs) are a platform of choice for gene therapy. However, liver-directed transduction has been hindered by immune responses unpredicted in the preclinical models, resulting in therapy failure. Liver immune responses are strictly regulated by the interactions between hepatocytes and non-parenchymal liver cells, such as Kupffer cells (the liver-resident macrophages) but how rAAVs induce such responses remains largely unknown. Therefore, human models recapitulating such interactions are required to address innate immune responses. Here, we developed a human 3D model to characterize the contribution of hepatocytes and Kupffer cells to the innate immune response. We developed a strategy for the differentiation of Kupffer-like cells from circulating monocytes based on cell-cell contact with primary human hepatocytes. We fine-tuned critical co-culture parameters to obtain a Kupffer-like phenotype while retaining hepatocyte viability and identity. Functional assessment of the differentiated 3D co-cultures showed that the model is responsive to classical pathogen-associated molecular pattern molecules, at a gene expression and secretory level. Moreover, we observed increased proinflammatory cytokine expression and secretion when challenged with a rAAV vector. Our data indicate the suitability of the novel model to investigate hepatocyte-Kupffer cell interactions and address innate immune responses within a human liver microenvironment.

## Introduction

The liver is a heterogeneous organ that exerts a crucial role in detoxification, metabolism, and production of essential proteins[1]. It is characterized by a tolerogenic phenotype due to its connection to the gastrointestinal tract via the portal vein, which delivers a continuous influx of non-self antigens to the hepatic immune environment[2]. The tightly regulated interactions between parenchymal and non-parenchymal cells (NPCs) are fundamental to maintain organ homeostasis and prevent excessive inflammation[3], [4]. Kupffer cells (KCs), the liver-resident macrophages, play a central role in such mechanisms through the production of anti-inflammatory cytokines, such as IL-10. However, upon stimulation they can release a variety of pro-inflammatory cytokines and chemokines (i.e., IL-6, IL-1β, TNF-α, IL-8) and initiate innate immune responses, representing the first line of defense against pathogens[5], [6], [7], [8]. Additionally, both hepatocytes and other NPCs, such as liver sinusoidal endothelial cells, can also play a role in mediating such responses by further activating KC, recruiting neutrophils and monocytes to the liver, and resolving the inflammation process[9]. Moreover, both parenchymal and non-parenchymal liver cells can exert a role as non-classical antigen-presenting cells (APCs), as they express a variety of pattern recognition receptors (PRRs) and class I and II major histocompatibility complex (MHC) molecules that can lead to T-cell activation[10], [11], [12], [13], [14]. However, how these cells interact to regulate innate immune processes is not fully elucidated. Although animal models provide valuable insights into the mechanisms regulating this intricated crosstalk, they fail to reproduce human disease pathogenesis or predict drug responses[15]. The failure in depicting such intricate interactions led to severe limitations in the development of new therapies. In the context of gene therapy, the inadequacy of pre-clinical models in predicting innate immune responses in the liver resulted in the development of CD8+ T-cell-mediated cytotoxic responses during clinical trials employing recombinant adeno-associated viruses (rAAVs), ultimately leading to therapy failure[16], [17]. This is not surprising as cross-species comparisons individuated differences in immune responses to different types of stimuli[15], [18], [19], highlighting the limitations that these models impose in a pre-clinical setting. Hence, human-based *in vitro* cell models are needed to unravel the liver cell interactions that regulate innate immune responses. While cell lines offer easy manipulation and reproducibility, their lack or low expression levels of sensing pathways compared to primary cells hamper their use in innate immune response evaluation[20], [21]. As such, primary human hepatocytes (PHHs) represent the gold standard for pre-clinical assessment, although difficulties in the maintenance of their function over extended periods have been reported[22]. However, it is well established that 3D primary hepatic cell cultures exhibit higher long-term viability, metabolic activities, and cell functions compared to 2D cultures[23], [24], [25]. On the other hand, the high costs and the scarce availability of primary materials, especially for the NPC compartment, have limited the development and employment of fully human *in vitro* 3D preclinical models.

Herein, we aimed to develop a primary human cell-based 3D model to characterize the contribution of hepatocytes and KCs to the innate immune response. Our group previously developed a method for culturing PHHs in 3D, in perfusion stirred-tank bioreactors, which allows the maintenance of their phenotype and functionality for extended periods[26], [27]. We have recently adapted the method to a smaller scale, overcoming limitations imposed by the scarcity of PHHs[28]. Here, we leveraged such PHH 3D culture to develop a strategy to differentiate KCs from peripheral blood mononuclear cell (PBMC)-derived monocytes, circumventing the limited availability of primary KCs. We hypothesized that by co-culturing PBMC-derived monocytes with PHHs, we would recapitulate KC differentiation, mimicking the KC niche replenishment via recruitment and differentiation of circulating monocytes happening upon liver infection or injury[29], [30]. Therefore, we developed a protocol to attain efficient differentiation into functional Kupffer-like cells (KLCs), while retaining high PHH viability and identity. The data presented herein show that our primary cells-based PHH:KLC co-culture is suitable for the assessment of innate immune responses within a human-based liver microenvironment, leveraging further studies for a deeper understanding of liver parenchymal and NPC crosstalk.

## Results

### Establishment of a small-scale 3D hepatic model employing primary human hepatocytes (PHHs)

We set to establish a robust KLC differentiation strategy to develop a 3D co-culture of PHHs and KLCs and evaluate their combined contributions to the development of an innate immune response within a human liver microenvironment. We leveraged from a protocol previously established by our team for the long-term culture of PHH with HepaRG cells in 30 mL stirred-tank vessels[28]. We adapted such protocol to obtain 3D PHH monocultures, that could be further co-cultured with KLC, while maintaining hepatic identity and function (Fig. 1). Briefly, PHHs were inoculated as a single cell suspension and maintained under agitation (80 rpm) for seven days, with a 20% media replacement every day after the third day of 3D culture (Fig. 1a). After 7 days, spheroids of different PHH lots retained a compact morphology and cell viability, as corroborated by the detection of few scattered propidium iodide (PI)-positive nuclei (Fig. 1b). The accumulation of F-actin throughout the apical region of the cells suggested that cell polarization, typical of hepatocytes, was preserved in the 3D culture. Hepatocyte identity was confirmed by the detection of the liver-specific hepatic nuclear factor 4 alpha (HNF4-α), albumin synthesis, and the expression of the major intermediate filament protein of the liver, cytokeratin-18 (Fig. 1c), together with the expression of *HNFA, ALB* and *CYP3A4* at gene level (Fig. 1d). Altogether, our results confirm the successful adaptation of our previous co-culture protocol for the monoculture of PHH spheroids while maintaining liver phenotype.

**Figure 1.**
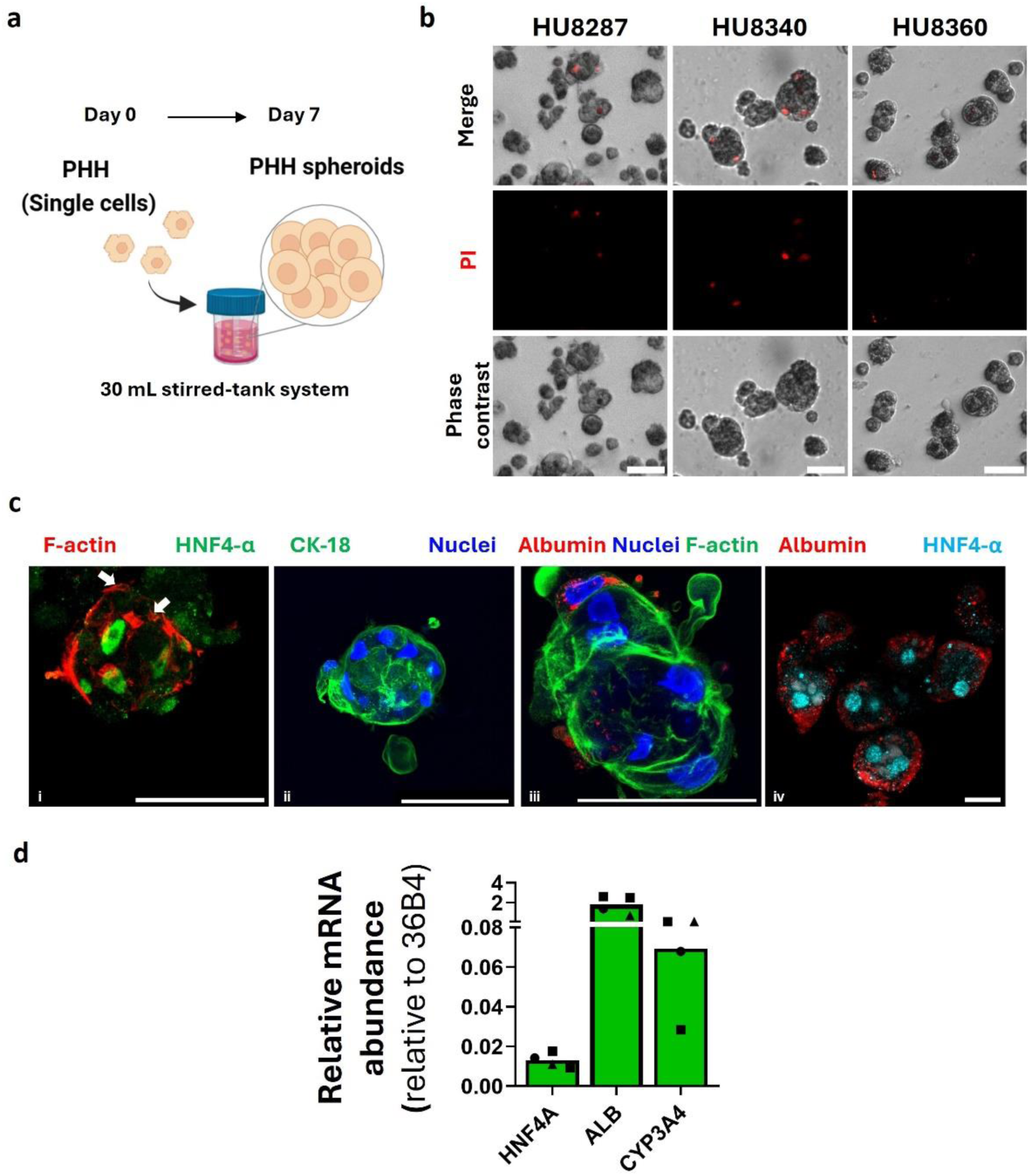
PHH spheroid cultures retain hepatocyte viability, morphology and phenotype. Schematic representation of PHH aggregation strategy in 30mL stirred-tank system. B) Aggregation and cell death evaluation of three different PHH lots at day 7 of 3D culture. Cell death assessed by staining with PI (red); scale bar = 50 µm. C) Evaluation of liver-specific markers in PHH spheroids at day 7 of 3D culture of different PHH donors (representative pictures). Immunofluorescence confocal microscopy of (i) F-actin (phalloidin, red), HNF4-α (green), (ii) Cytokeratin-18 (green), (iii) Albumin (red), F-actin (phalloidin, green), (iv) Albumin (red), HNF4-α (light blue). Nuclei were stained with DAPI (blue) in (ii) and (iii). The white arrows point to possible bile canaliculi-like structures. Image analysis was performed with imageJ Maximum Intensity Z-projection tool (5/6 stacks per image, 1 μm stacks); scale bars = 50 µm. D) Gene expression evaluation of hepatic markers in 3D PHH (7 days of 3D culture). Data are expressed as relative mRNA abundance relative to 36B4; N = 4. Different symbols correspond to different PHH donors.

### Impact of liver microenvironmental cues on KLC differentiation

To obtain KLCs, we assessed how PHHs and liver microenvironmental cues could impact the differentiation of PBMC-derived monocytes, mimicking monocyte recruitment to the liver upon injury[29], [30], [31]. We optimized medium composition to maintain both hepatic and myeloid identity and function. As such, the effect of fresh PHH medium and 3D PHH-conditioned medium was assessed on monocytes isolated from PBMCs (Supplementary Fig. S1). We also tested the effect of dexamethasone on KLC differentiation, as dexamethasone has been shown to have a mitigating impact on immune responses[32], [33], but its role in hepatocyte maturation and maintenance is well established[34]. After 6 days of culture in PHH medium with dexamethasone, cells acquired a rounded morphology, while a spindle-like phenotype was observed in unstimulated macrophages (M0-like) and macrophages cultured in PHH medium without dexamethasone (Supplementary Fig. S1a). Moreover, PHH medium supplemented with dexamethasone (fresh or 3D PHH-conditioned) induced an increase in the expression of a curated panel of KC markers [35], [36] (Supplementary Fig. S1). Importantly, we observed a significant upregulation in *MARCO*, reported to be unique to the resident subset of the human liver macrophage population[35]. Altogether, our results indicate that dexamethasone is beneficial for the differentiation of the KLC population. Of note, none of the tested PHH media induced a pro-inflammatory phenotype in the differentiated myeloid population, as evidenced by a significantly lower expression of *Dectin2* in comparison to pro-inflammatory macrophages [37] (Supplementary Fig. S1b).

To evaluate the effect of cell-cell interactions on the differentiation of the KLC population, we co-cultured PHH spheroids and PBMC-derived monocytes in a 1 PHH:2 monocyte ratio and assessed the expression of the KC marker panel after 6 days of co-culture (Fig. 2a). Moreover, we fine-tuned media composition by evaluating the effect of CCL2 and M-CSF supplementation on KLC differentiation efficiency, given their roles in monocyte recruitment and macrophage maintenance during inflammation processes and homeostasis[31], [38], [39]. We observed KC marker expression when monocytes were co-cultured with PHH spheroids for 6 days (Fig. 2a, left panel), confirming the integration of the myeloid population in the co-culture. Importantly, the hepatic marker expression was maintained in the co-culture (Fig. 2a, right panel), indicating that neither the presence of the myeloid population nor its differentiation process affected the hepatic identity. Although no significant differences were observed between conditions, a tendency for a higher expression of KC markers was detected both at gene and protein levels when supplementation was applied (Fig. 2a, left panel, Fig. 2b, Supplementary Fig. S2, top panel). Moreover, supplementation did not affect the activation status of the myeloid population or the expression of hepatic markers in the co-culture (Supplementary Fig. S2, middle/bottom panels). Overall, these results point to a non-detrimental effect of MCSF and CCL2 supplementation, with a possible role for the achievement of an efficient differentiation process.

**Figure 2.**
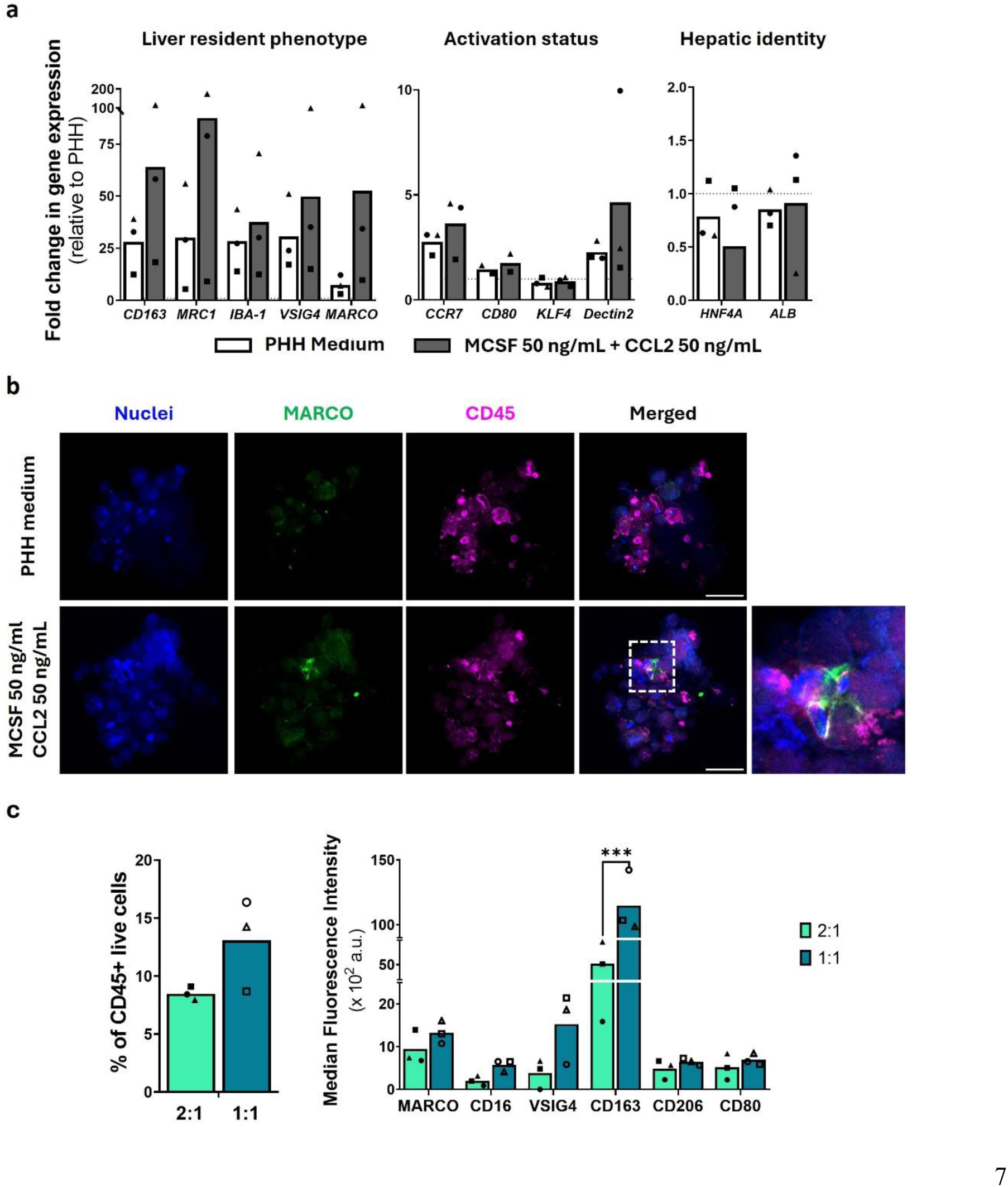
Supplement addition and ratio optimization maintain the phenotype of the KLC and PHH populations in the 3D co-cultures. A) Gene expression analysis of markers for Kupffer cell identity (left graph), activation status (central graph) and hepatic identity (right graph) of day 6 static 3D co-cultures with or without M-CSF 50 ng/mL and CCL2 50 ng/mL supplementation. Data are normalized for a housekeeping gene expression (36B4) and expressed as fold change in respect to 3D PHH cultures. N = 3 (CD80 N=2). B) Assessment by confocal microscopy of the differentiated myeloid population in 3D PHH:KLC static co-cultures in the two different media compositions for 6 days. Co-cultures were stained for CD45 (Pink) and MARCO (Green), nuclei were stained with DAPI (blue). Image analysis performed with imageJ Maximum Intensity Z-projection tool (5/6 stacks per image, 1 μm stacks); scale bars = 50 µm. C) Myeloid population maintenance and Kupffer cell phenotype evaluation in 3D dynamic co-culture strategy employing a 1:1 or 2:1 PHH:monocyte ratio. Results are expressed as percentage of CD45-positive cells (left graph) and each marker Median Fluorescence Intensity within the CD45-positive population (right graph) after 6 days of co-culture . N = 3. Two-way ANOVA was performed for statistical analysis. The same PHH donor was employed for media and ratio optimization. In all graphs, symbols correspond to different PHH:KLC combinations.

Lastly, we optimized the cell ratio aiming to maximize the differentiation of monocytes into KLCs by increasing hepatocyte-myeloid cell interactions. Therefore, we tested under dynamic conditions a 1 PHH:1 monocyte ratio and compared KC differentiation efficiency in respect to the 1:2 ratio employed for the previous optimization steps. Both cell ratios led to the detection of a CD45+ population characterized by the expression of the KC marker panel after 6 days of co-culture (Fig. 2c). However, the employment of a larger fraction of monocytes at the beginning of the differentiation process resulted in a higher percentage of myeloid cells at day 7. Moreover, a significant difference in CD163 expression and an overall tendency for higher expression of KC markers was observed when employing the 1:1 ratio, without altering the expression of activation markers (CD80).

### Characterization of PHH:KLC 3D dynamic co-cultures

Following the individuation of critical co-culture parameters (media composition, cell ratio), we aimed to establish a robust protocol for the 3D differentiation and co-culture of PHHs and KLCs in a dynamic system (Fig. 3a). To guarantee the reproducibility of the process, we employed 3 different PHH donors in combination with monocytes isolated from 5 different PBMC healthy donors and assessed differences in the co-culture outcome. Independently of the combination employed, monocyte addition to PHH spheroids in the 30 mL stirred-tank system did not affect hepatic aggregation, with no differences in viability at 13 days respect to 7 days of 3D culture (Fig. 3b, Supplementary Fig. S3a). Flow cytometry analysis showed that the selected co-culture strategy led to the expression of KC markers in the CD45+ population, with a significant increase in the percentage of positive cells for CD16, MARCO, VSIG4 and CD163 in respect to the initial monocyte population (Fig. 3c, Supplementary Fig. S3b). Remarkably, differentiation into KLC was observed regardless of the PHH or PBMC donor employed, highlighting the robustness of the differentiation process.

**Figure 3.**
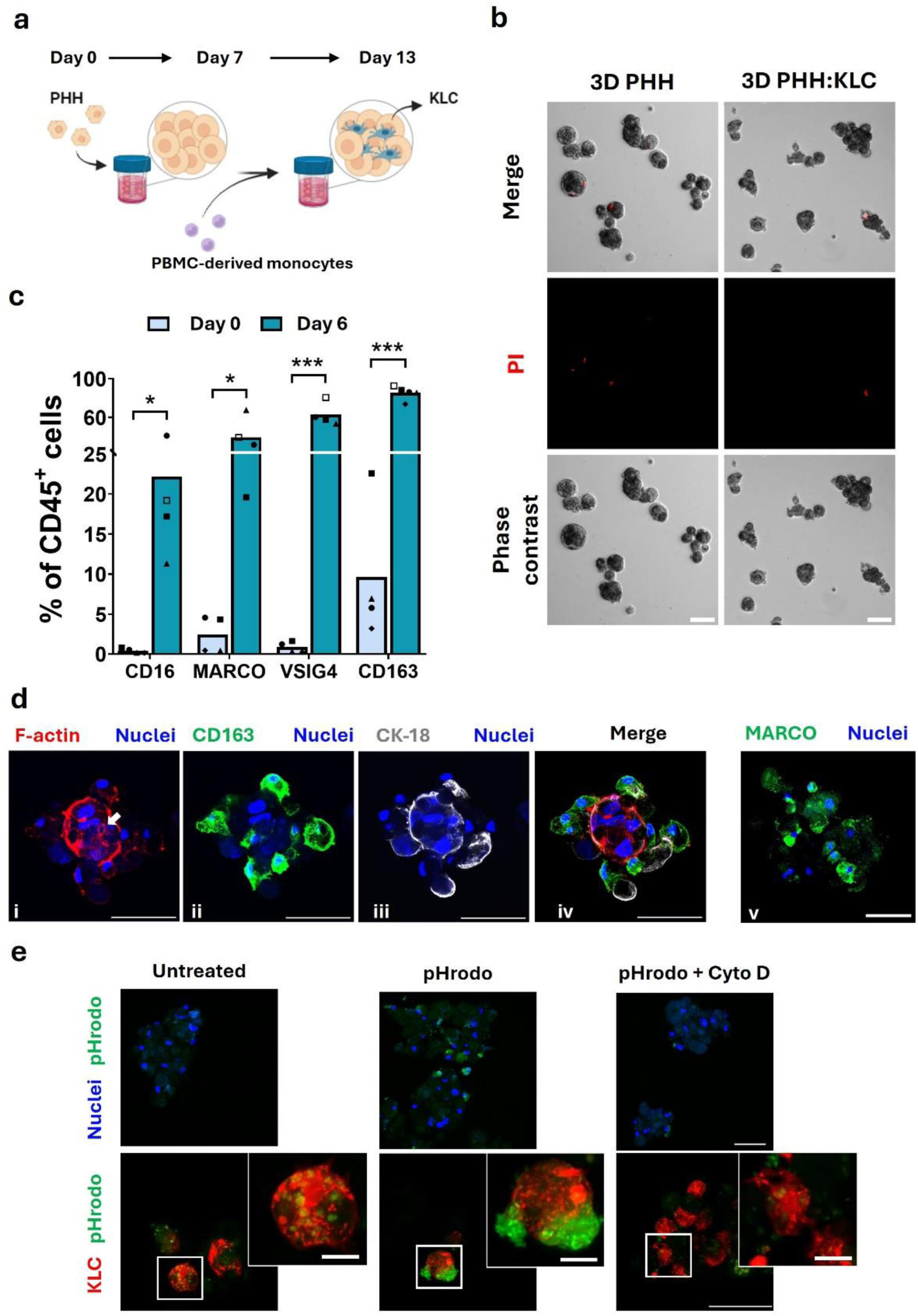
PHH:KLC spheroid cultures retain viability and a Kupffer-like phenotype while maintaining hepatic and macrophage functionality. A) Schematic representation of PHH:KLC aggregation strategy in 30 mL stirred-tank system. B) Aggregation and cell death evaluation of PHH (left panel) PHH:KLC (right panel) dynamic co-cultures at day 13 of 3D culture (13 days for PHH monoculture, 7 of PHH monoculture + 6 days of co-culture for PHH:KLC co-culture). Cell death assessed by staining with PI (red); scale bar = 50 µm. C) Flow cytometry evaluation of Kupffer cell markers in monocyte (Day 0) and KLC cells (Day 6 of co-culture). Results are expressed as percentage of positive cells detected inside of the CD45-positive population. Symbols correspond to different PHH:KLC combinations; N ≥ 4. Unpaired T test was performed for statistical analysis. D) Immunofluorescence microscopy of PHH:KLC 3D co-cultures at day 13 of 3D culture. Spheroids were stained for (i) F-actin (phalloidin, red), (ii) CD163 (green), (iii) Cytokeratin-18 (white) and (v) MARCO (green). Nuclei were stained with DAPI (blue). The white arrow points to apical F-actin accumulation. Image analysis was performed with imageJ Maximum Intensity Z-projection tool (3 stacks for image i, ii, iii and v, 10 stacks for image iv (merge), 1 μm stacks); scale bars = 50 µm. E) Phagocytic activity evaluation of PHH:KLC co-culture by pHrodo^TM^ assay. Samples were incubated with pHrodo^TM^ green BioParticles^TM^ (250 μg/mL) for 6h with or without pre-incubation with 10 μM Cytochalasin D (Cyto D, phagocytosis inhibitor) for 3h. In the top panel PHH:KLC spheroid were stained with DAPI (blue), in the bottom panel KLC were intracellularly labelled with cell Tracker deep red (red). 3-5 stack images, 1-2 µm stacks. Scale bars = 50 µm, zoom insert scale bars = 10 µm.

Immunofluorescence confocal microscopy analysis confirmed that the myeloid population was differentiated (CD163, MARCO) and integrated into the PHH-spheroid without altering hepatocyte identity (Fig. 3d). This was evidenced by the apical accumulation of F-actin (indicated with a white arrow) and CK-18 detection (Fig. 3d). Gene expression analysis further confirmed these results, with no differences detected in HNF4α, Albumin and CYP3A4 gene expression between 3D PHH monoculture and PHH:KLC co-culture (Supplementary Fig. S4a).

To assess the functionality of the differentiated KLCs, we tested their ability to perform phagocytosis, one of the main functions of macrophages[8]. We employed pHrodo™ Green E. coli BioParticles™ coupled with a conjugate that is non-fluorescent outside the cell but emits fluorescence when in contact with the acidic environment of the phagosome. When co-cultures were treated with the bioparticles for 6h, pHrodo™ fluorescence was observed, with internalized particle detection corresponding to the myeloid cells (Fig. 3e), as also observed in 2D M0 macrophages (Supplementary Fig. S4b). Fluorescence was not detected when co-cultures were pre-treated with a phagocytosis inhibitor (Cytochalasin D) (Fig. 3e) or in PHH monocultures with any of the tested conditions (Supplementary Fig. S4c), indicating the specificity of the assay for the myeloid population.

Hence, our results indicate that the optimization of critical co-culture parameters led to the definition of a robust KLC differentiation protocol. Moreover, we were able to implement a 3D co-culture system in which PHHs maintain hepatocyte identity and the myeloid population is characterized by a functional KLC phenotype, suitable for the evaluation of PHH-KC interactions.

### PHH:KLC 3D co-culture are responsive to inflammatory stimuli

To evaluate the ability of the 3D cultures to respond to stimuli, we activated different Toll-like receptor (TLR) downstream pathways by challenging the co-culture with prototypical Pathogen Associated Molecular Patterns (PAMPs), namely Pam3CSK4 (TLR2 agonist) and lipopolysaccharide (LPS, TLR4 agonist). Moreover, we tested the ability of the culture to respond to genomic DNA through stimulation with ADU-S100, an agonist of the Stimulator of Interferon Gene (STING) pathway, which plays a central role in antiviral innate immunity[40]. After 6h of challenge, we evaluated the expression of key downstream cytokine genes (coding for IL-8, IL-1β, IL-6, and TNFα) to assess the 3D cultures responsiveness to each type of stimulus (Fig. 4, Supplementary Fig. S5). In response to LPS, both PHH and PHH:KLC cultures upregulated the expression of the genes *CXCL8*, *IL1B*, *IL6* and *TNFA* (Fig. 4a). Both cultures showed an increase in gene expression in response to TLR2 agonist, with a higher induction observed in the co-culture with a significant upregulation of *CXCL8, IL1B, IL6* and *TLR2* (Fig. 4b). These results were further confirmed by the detection of secreted IL-6, a main cytokine secreted by PHHs during acute phase inflammation[13], [41]. A trend for IL-6 secretion was detected as early as 6h of challenge with LPS in both PHH and PHH:KLC cultures, while IL-6 secretion increase was only observed in the presence of the myeloid population in the case of Pam3CSK4 challenge (Fig. 4c). The co-culture also demonstrated to be responsive to STING stimulation, with a significant difference in *IL-6* expression, while no response was detected in PHH monocultures (Supplementary Fig. S5a), in agreement with previous reports in human and murine hepatocytes[42]. Interestingly, a different level of each Pattern Recognition Receptor (PRR) and relative downstream cytokine gene expression was observed between unstimulated PHH and PHH:KLC cultures, with a significant difference in *CXCL8, TNFA* and *STING* basal expression and an overall higher receptor availability and cytokine gene expression in the co-culture (Supplementary Fig. S5b-c) that could have led to a prompter response in respect to the PHH monoculture. Altogether, our results confirm the functionality and suitability of our 3D co-culture model for the assessment of innate immune responses.

**Figure 4.**
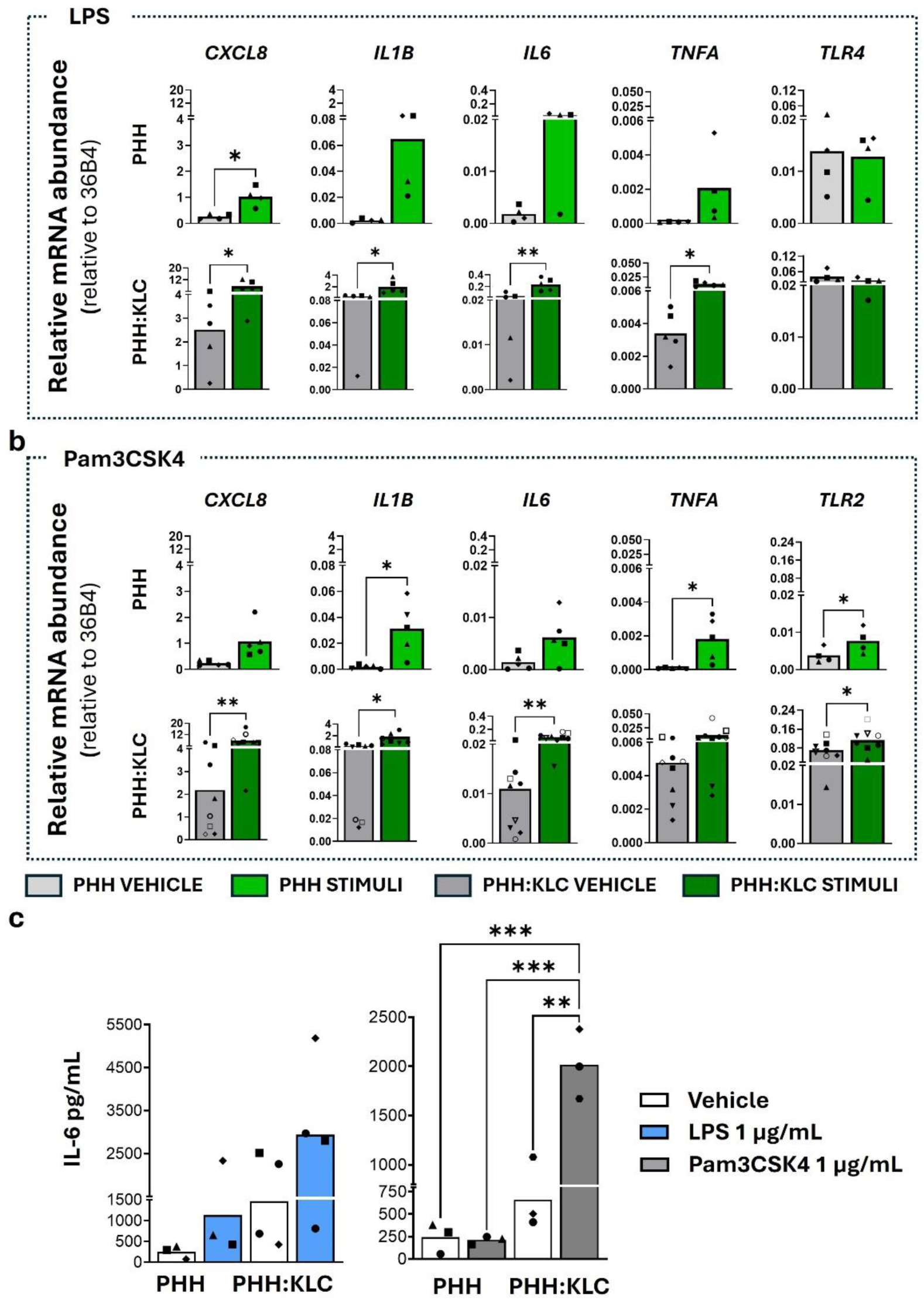
3D PHH and 3D PHH:KLC cultures are responsive to PAMPs. Gene expression evaluation of PRRs and cytokines after stimulation with prototypical stimuli for 6h in PHH and PHH:KLC (13 days of 3D culture). Cultures were stimulated with: A) LPS 1 µg/mL, B) Pam3CSK4 1 µg/mL. Data are expressed as relative mRNA abundance relative to 36B4; N ≥ 3. Paired T-test was performed for statistical analysis. C) Evaluation of IL-6 secretion in PHH and PHH:KLC 3D cultures after stimulation with LPS 1 µg/mL or Pam3CSK4 1 µg/mL; N=3. One way ANOVA was employed for statistical analysis. In all graphs, different symbols correspond to different PHH:KLC donor combinations.

### IL6 is upregulated in PHH:KLC 3D co-cultures challenged with rAAV

To demonstrate the applicability of our co-culture system for the evaluation of rAAV-driven innate immune responses, we firstly assessed the cellular tropism of an rAAV8 encoding mCherry under the control of the cytomegalovirus (CMV) promoter. Briefly, we exposed both PHH and PHH:KLC 3D co-cultures to the viral vector employing an MOI of 1x10^6^ VG/cell. As soon as 6h after exposure to the viral vector, we could detect transgene expression in the co-cultures by ddPCR, which increased at 24h in both PHH and PHH:KLC 3D cultures (Fig. 5a, Supplementary Fig. S6a). Immunofluorescence analysis of the co-cultures transduced with the viral vector for 7 days confirmed transgene expression, with mCherry detection observed in PHH but not in KLC (Fig. 5b). Accordingly, transgene expression was observed in PHH monocultures both in 3D and 2D, while no expression was observed in M0-like macrophages (Supplementary Fig. S6b-d), suggesting a preferential transduction of the virus in PHH.

**Figure 5.**
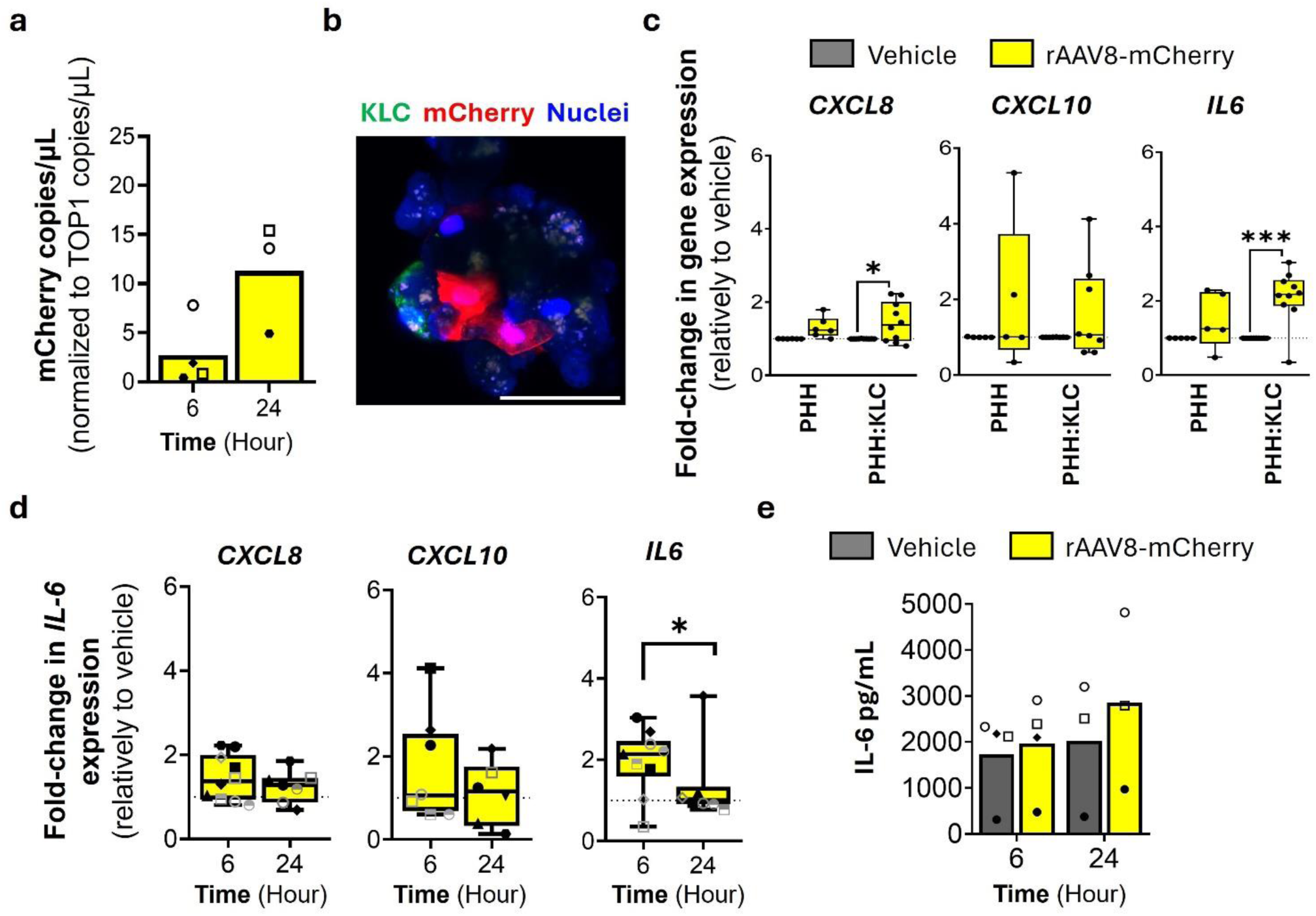
3D PHH:KLC cultures are susceptible and responsive to rAAV8. A) Evaluation of mCherry gene expression after PHH:KLC 3D cultures transduction with rAAV8-mCherry for 6 and 24h. Results are expressed as fold change in respect to the vehicle condition; N = 3. B) Immunofluorescence confocal microscopy evaluation of PHH:KLC 3D co-cultures transduced with rAAV8-mCherry (MOI 1x10^6^ VG/cell). Myeloid cells were intracellularly labelled with Cell Tracker deep red (green) and nuclei were stained with DAPI (blue). Transgene expression was assessed by mCherry detection (red); 1 stack image, scale bar = 50 µm. C) Gene expression evaluation of cytokines after stimulation with rAAV8-mCherry for 6h in PHH and PHH:KLC (13 days of 3D culture). Results are normalized for a housekeeping gene expression (36B4) and expressed as fold change in respect to the vehicle condition; N ≥ 8. Unpaired T-test was performed for statistical analysis. D) Gene expression evaluation of cytokines after stimulation with rAAV8-mCherry for 6 and 24Hh in PHH:KLC (13 days of 3D culture). Results are normalized for a housekeeping gene expression (36B4) and expressed as fold change in respect to the vehicle condition; N ≥ 6. Paired T-test was performed for statistical analysis. E) IL-6 secretion evaluation after PHH:KLC 3D cultures challenge with rAAV8-mCherry for 6 and 24h. Paired T-test was performed for statistical analysis; N ≥ 3. In all graphs, symbols correspond to different PHH:KLC donor combinations. For box plots, the whiskers denote minimum and maximum values and the horizontal line the median.

We then evaluated the ability of the co-culture to mount an immune response against the viral vector. We assessed *IL-6, CXCL8* and *CXCL10* gene expression after 6, and 24h of challenge with rAAV8-mCherry given their implication in viral sensing responses[43]. At 6h post-transduction, we detected a significant increase in *IL6* and *CXCL8* expression relative to the control condition (Fig. 5c). Such response was not observed in PHH monocultures, highlighting the necessity of the myeloid population to recapitulate rAAV-mediated innate immune responses in the liver. From 6h to 24h post-transduction, we observed a decrease in expression of all tested cytokines (Fig. 5d), in line with the fast and transient nature of the innate immune response. Notably, our co-culture system successfully captured it, despite the variability introduced by the employment of primary material from different donors. Accordingly, IL-6 showed a trend for increased secretion when compared to the vehicle, at both 6 and 24h (Fig. 5e). Nevertheless, the donor-to-donor variability observed affected the magnitude of such increase in each tested culture. Overall, these results suggest that the co-culture developed herein could be employed to leverage further studies aimed at tackling the rAAV-innate immune responses within a human liver microenvironment.

## Discussion

In this study, we report the successful differentiation of PBMC-derived monocytes into Kupffer-like cells in a 3D human liver *in vitro* model. The strategy allows for the evaluation of hepatocyte and Kupffer cell-driven innate immune responses within a human liver context. This small-scale co-culture system overcomes the limitations of primary material availability while allowing for a uniform production of PHH:KLC spheroids. This methodology can thus be employed for the simultaneous evaluation of different types of immune stimuli and further investigate the cellular crosstalk between KC and PHH.

Most of the current preclinical models for the study of liver innate and adaptive immune responses still rely on animals, despite the reported limitations in the correlation with human immune responses18. On the other hand, human 2D cell cultures fail to recapitulate important aspects of structural, mechanical, and biochemical cues that dictate native cell interactions[25], not allowing for a suitable representation of the cellular crosstalk that regulates innate immune responses. Therefore, 3D human-based *in vitro* models are required for the elucidation of the mechanisms driving the immune response in patients. However, the correlation of such preclinical models with the clinical outcome is nevertheless dependent on the selected cell source. Stem cell-derived models have been developed to mimic liver inflammation and could represent a valuable tool to overcome the scarcity of primary cell sources[44], [45]. However, the lack of standardization of both parenchymal and non-parenchymal cell differentiation protocols, and the resulting maturity and functionality of the differentiated cells, hamper their use in preclinical settings. As such, primary cultures still constitute the gold standard for drug metabolism and immune response evaluation *in vitro*. As primary 3D liver cultures present higher long-term viability and cell functions compared to 2D cultures[23], [24], [25], they currently represent the most reliable model to mimic the liver microenvironment *in vitro*[46]. Three-dimensional PHH cultures have been previously established by our group in 200 mL stirred-tank bioreactors (STB), maintaining for over a month in culture hepatic cell polarization, biosynthetic functions and xenobiotic metabolism[26], [27]. Recently, we have reduced the scale to 30 mL spinner vessels for the co-culture of PHH and HepaRG cells, allowing for the use of fewer primary hepatocytes[28]. Herein, we adapted such protocol to obtain a 3D PHH monoculture while maintaining the compact morphology and high cell viability observed in the previous systems (Fig. 1). The accumulation of F-actin in intercellular junctions suggests the correct repolarization of PHHs[47]. Moreover, the expression of HNF4-α, crucial transcription factor in mature hepatocytes[48], and albumin, one of the liver’s main secreted proteins[49], corroborate the suitability of the culture to address hepatocyte functions.

In healthy livers, tissue-resident macrophages arise from yolk-sac-derived erythro-myeloid progenitors[50]. However, upon injury or strong inflammation processes Kupffer cells can be depleted and the liver gets repopulated by recruiting bone marrow-derived circulating monocytes. These monocytes gradually lose expression of monocytic genes while acquiring Kupffer cell characteristics[29], [30], [31]. We therefore hypothesized that co-culturing human monocytes isolated from peripheral blood mononuclear cells with hepatocytes would recapitulate KC differentiation by mimicking monocyte recruitment and differentiation *in vivo*. Our data show that the novel differentiation strategy robustly led to a myeloid population with typical KC marker expression and functions (Fig. 3), in agreement with reports of KC differentiation from bone marrow-derived monocytes in mice[31], [36]. An increase in KC markers was always observed, regardless of the PHH or PBMC donor employed (Fig. 3C), underlying that the donor mismatch between the two cell sources does not compromise the efficient differentiation of the myeloid population. Importantly, the introduction of monocytes in the 3D PHH monoculture did not affect the hepatocyte phenotype (Fig. 2a). We therefore report on the development of a fully human 3D model based on primary cells that can be employed to tackle both hepatocyte and KC functions within a liver microenvironment. Importantly, the employment of a small-scale stirred-tank system and the developed differentiation strategy allows for circumventing the limitations of the scarce primary material availability for both cell types.

Upon PAMPs recognition, the different TLRs undergo oligomerization and recruit adaptor molecules to engage downstream signaling proteins[51]. Among them, TLR4 signaling in the liver is one of the best characterized, mediated by both parenchymal and non-parenchymal cells[52], [53]. Through both myeloid differentiation primary response 88 (MyD88)-dependent or MyD88-independent signaling pathways, the transcription factors nuclear factor (NF)-κB and activator protein (AP-1) or the interferon regulatory factor (IRF) 3 can be activated, leading to increased expression of a variety of pro-inflammatory cytokines, type I interferons and interferon-responsive genes[51], [52]. Our data showed a clear induction of pro-inflammatory cytokine expression by the co-culture system upon LPS stimulation (Fig. 4a), confirming the functionality of the differentiated myeloid population and the suitability of the model for the assessment of TLR4-mediated liver innate immune responses. The 3D PHH monoculture analysis also showed an increase in cytokine expression in response to LPS, in agreement with reports of IL-6 induction against this PAMP in both animal and human models[6], [32], [41]. However, the response differs between the two 3D culture systems, highlighting the influence of PHH-KC crosstalk in the induction of liver innate immune responses. This is even more evident when challenging cultures with the TLR2 agonist Pam3CSK4, where higher cytokine gene expression was observed in the co-culture (Fig. 4b), or with the STING agonist ADU-S100, where *IL6* expression increase was observed only in the co-culture system (Supplementary Fig. S5a). Given that the co-culture of PHH with KLC did not lead to evident changes in expression of typical activation genes (CCR7, CD80, KLF4, Dectin2) when compared to PHH monocultures (Fig. 2a), such differences in immune response might be explained by the higher basal cytokine expression in the PHH:KLC 3D culture relatively to the PHH monoculture (Supplementary Fig. S5b-c). A higher cytokine availability could have led to a faster and greater activation of the different innate immune pathways, resulting in differences in the kinetic and magnitude of response between the two 3D cultures. Importantly, we observed response to pro-inflammatory stimuli despite the presence of dexamethasone in the co-culture medium, which is reported to decrease immune cell response[32], [33]. An *in vitro* 3D human PHH:KC model showed that concentrations of 50 nM or lower of dexamethasone allow for increased IL-6 secretion in response to 10 μg/mL LPS for 48h[32]. Accordingly, the concentration selected for our co-culture (18.75 nM) enabled the detection of cytokine increase at both gene and protein levels as early as 6h post-stimulation. Altogether, these results show that the PHH:KLC co-culture is responsive to various prototypical PAMPs and can be employed to unravel innate immune responses within a human liver microenvironment. Further characterization is required to understand how the PHH-KLC crosstalk influences the response through the different sensing pathways.

rAAVs are vectors of choice for gene therapy. However, despite their expected low immunogenicity, liver-directed transduction with rAAVs has been hindered by unpredicted immune responses mediated by neutralizing antibodies (humoral immunity) and CD8+ T-cells (adaptive immunity)[16], [17]. Animal models have been useful tools in addressing the humoral response but have failed to reproduce the adaptive immunity induced by rAAVs. As a result, the innate immune responses mediated by rAAVs and the respective pathways are yet to be elucidated in the human liver. In mice, KCs were shown to respond to double-stranded DNA (self-complementary rAAVs), mainly via TLR9, with subsequent activation of CD8+ T-cells[54], [55]. A report employing a human co-culture liver model suggested a TLR2-mediated response against rAAVs by KCs and other NPCs, but not by hepatocytes[56]. More recently, studies employing human and non-human primate PBMCs *in vitro* demonstrated the secretion of IL-6 and IL-1β as a major response to rAAV capsids, with a reversion of the rAAV-driven immunotoxicity *in vivo* by the neutralization of their respective receptors[57], [58]. However, the scarce representation of the human liver immune cell crosstalk in such models hampered a deeper characterization of these responses[15], [25]. As a role for TLR2, TLR4 and STING pathways in viral sensing has been previously reported[40], [59], [60], [61], we hypothesized the applicability of our co-culture to evaluate rAAVs-induced innate immune responses. In agreement with previous reports, our results show a significant upregulation in *IL6* and *CXCL8* after 6h of transduction, and a tendency for a higher expression of *CXCL10* and IL-6 secretion (Fig. 5). Notably, our results corroborate previous studies implying IL-6 secretion as an early human immune response to rAAVs[58]. Cytokine expression was generally decreased after 24h of rAAV transduction, confirming the rapid nature of rAAV-induced innate immune responses. In fact, the generally mild increase in cytokine gene expression observed in our co-culture system aligns well with the transient nature of the rAAV-induced immune responses and the relatively low immunogenicity previously reported for these vectors[58], [62], leading to a generally low detection of rAAV-driven innate immune responses in the liver[58], [62], [63]. Of note, despite the consistent up-regulation of *IL6*, its peak of expression kinetics varied depending on the PBMC donor employed (Fig. 5d), highlighting the co-culture ability to detect donor-specific variations. Differences between the PBMC and/or PHH donors employed (i.e., previous AAV exposure and infection status at the time of collection) could have an impact on both kinetics and magnitude of rAAV-driven innate immune responses, explaining the observed variability among the different cultures. Further studies are required to thoroughly characterize the specificities of the dynamic and sensing pathways that lead to the development of the adaptive responses observed in patients. The 3D co-culture could also be leveraged to include additional NPCs, allowing to foster further cell-cell interactions contributing to the liver innate immune response[9], hence increasing the physiological relevance of the model. Nonetheless, our data show that the herein presented PHH:KLC co-culture system can detect increased cytokine expression and distinguish donor-to-donor differences, pointing to the applicability of this minimal cell configuration to address the rAAV-mediated mechanisms driving the immune responses observed in clinical trials.

Altogether, we report the successful development of a robust strategy for the differentiation of KLCs from PBMC-derived monocytes based on liver microenvironmental cues and contact with PHHs in 3D. The resulting primary-based human 3D model allows for the evaluation of both hepatocyte and KC-driven innate immune responses in a small and high throughput scale, which can be used to further investigate the cellular crosstalk between KCs and PHHs.

## Materials and Methods

### Human samples

Cryopreserved PHH lots were purchased from Thermo Fisher Scientific. Buffy coats from healthy donors were obtained from the Portuguese Institute for Blood and Transplantation (IPST, Lisbon, Portugal). Written informed consent was received from donors prior to the study (in conformity with IPM-74.52. V.4, IPST). Experiments employing human samples were approved by the Ethics Committee of the Institute of Hygiene and Tropical Medicine, the António Xavier Institute of Chemical and Biological Technology and the Faculty of Law of the Universidade Nova de Lisboa (CE IHMT-ITQB-NSL). All methods were performed following the Portuguese and European legislation and the commonly internationally accepted ethics directives and guidelines.

### HepaRG 2D monoculture

The HepaRG cell line was purchased from Thermo Fisher Scientific. Cells were thawed at 37°C and diluted 1:10 in William’s E medium, supplemented with 1 % (v/v) Glutamine, 1 % (v/v) penicillin/streptomycin, 10 % (v/v) fetal bovine serum (FBS) (all from Thermo Fisher Scientific), 5 μg/ml bovine insulin and 50 μM hydrocortisone hemisuccinate (both from Merck). Cells were plated at 2.6x10^4^ cell/cm^2^ in T75 flasks. Medium replacement was performed 24h after thawing and then twice a week until day 14 of culture, when cells were passaged. Conditioned medium from D14 of HepaRG culture was collected and centrifuged at 1000 xg, 5 min and stored at -20°C until further use in PHH 3D monocultures. HepaRG cells were passaged at least twice before conditioned media was collected. Cells were routinely tested for mycoplasma contamination.

### Primary human hepatocytes (PHH) 3D monocultures

PHHs lots (HU8287, HU8340, HU8360, HU8412) were thawed according to the vendor’s instructions. Briefly, vials were thawed at 37°C, diluted in 25 mL of cryopreserved hepatocytes recovery medium (CM7000, Thermo Fisher Scientific), and centrifuged at 100 xg, for 10 min. PHHs were resuspended in a 4:1 mixture of hepatocyte maintenance medium (PHH medium, Williams’ Medium E supplemented with Hepatocyte Maintenance Supplement Pack (Serum-free)) and 10% (v/v) Fetal bovine serum (FBS) (all Thermo Fisher Scientific) and HepaRG-conditioned culture medium. The cell suspension was inoculated in 30 mL spinner vessels (ABLE Biott Corporation®) at 5 x 10^5^ cell/mL to generate 3D PHH cultures. Spinner vessels were maintained under agitation (80-110 rpm) for 7 days. From day 3 until the end of the culture, a 20% medium exchange regimen with serum-free PHH medium was implemented, to remove daily both HepaRG-conditioned medium and FBS from the PHH 3D monocultures (to reach 0% HepaRG-conditioned medium and FBS at day 7 of 3D culture). PHH-conditioned medium was collected at D7 after the last media exchange, centrifuged at 1000 xg, 5 min and stored at -20°C until further use for 2D and 3D Kupffer cell differentiation.

### Peripheral blood mononuclear cell (PBMC) isolation

Buffy coats were collected and stored in blood collection bags at room temperature. Blood samples were slowly dispensed in a 50 mL falcon containing LymphoPrep (STEMCELL Technologies) without perturbing the liquid-liquid interface. Tubes were centrifuged at 950 xg, 25 min, without brakes. The PBMC layer was collected from the falcon tube and diluted in PBMC isolation buffer (PBS (-/-), 2% FBS, 2 mM EDTA). Samples were centrifuged at 400 xg, 10 min, washed with PBMC isolation buffer, and centrifuged at 400 xg, 10 min. The supernatant was discarded, and the pellet was resuspended in pre-warmed ACK lysing buffer (Gibco), for 5 min, at 37°C, to lyse any residual erythrocytes. Samples were diluted in PBMC isolation buffer and centrifuged at 400 xg, for 10 min. The resulting PBMCs were counted and stored in freezing medium (FBS with 10% DMSO), in the vapor phase of liquid Nitrogen.

### Monocyte isolation

Cryopreserved PBMCs were thawed in a ThawSTAR cell thawing equipment. Cells were gradually diluted in monocyte basal medium (RPMI 1640 with Phenol Red, 10% FBS and 1% P/S) and centrifuged at 400 xg, for 10 min. The supernatant was discarded, and the cell pellet was resuspended in monocyte isolation buffer (PBS (-/-), 2% FBS, 1 mM EDTA, 0.1 mg/mL of DNase I (STEM CELL Technologies) and incubated for 15 min at room temperature. The cell suspension was then filtered through a 40 μm strainer and cells were counted to assess PBMC recovery. Cells were resuspended at 50 X 10^6^ cell/mL and monocyte isolation was performed employing the EasySep human monocyte isolation kit (STEMCELL Technologies), following the manufacturer’s instructions.

### 2D monocyte differentiation

Freshly isolated monocytes were plated at a seeding density of 0.3 x 10^6^ cell/cm^2^ and maintained for 6 days in monocyte basal media supplemented with 50 ng/mL of M-CSF for M0-like macrophage differentiation or in PHH medium (with or without 0.1 μM dexamethasone or in a 1:1 mixture with conditioned medium from 7 days of PHH 3D culture) for 2D Kupffer-like cell differentiation optimization. At day 4 of differentiation, M0-like macrophage medium was exchanged, and cells were maintained for 2 additional days in monocyte basal medium supplemented with 10 ng/mL LPS and 50 ng/mL IFN-γ for pro-inflammatory (M1-like) macrophage differentiation or 20 ng/mL IL-4 and 20 ng/mL IL-13 for anti-inflammatory (M2-like) differentiation.

### 3D PHH viability and cell concentration assessments

PHH 3D culture viability was assessed by fluorescence microscopy using 1 µg/mL propidium iodide (PI; Merck) to label dead cells. Cell concentration was determined by digesting a sample of PHH spheroids with 0.05% Trypsin/EDTA (Thermo Fisher Scientific) and the resulting single-cell suspension was counted using a Fuchs-Rosenthal chamber by the trypan blue exclusion method.

### Static 3D co-culture

D7 PHH spheroids were centrifuged at 100 xg, for 1 min, and the supernatant was collected and centrifuged 1000 xg, for 5 min to remove debris. PHH spheroids were plated in flat ultralow attachment 96-well plates at 2.5 x 10^4^ cell/well, in 100 µL of PHH-conditioned medium. Isolated monocytes were added to the 3D culture in a ratio of 1 monocyte to 2 PHH (1.25 x 10^4^ cell/well) in fresh PHH medium without dexamethasone (final dexamethasone concentration of 50 nM). A 50 ng/mL Macrophage Stimulating Factor (M-CSF) and 50 ng/mL Monocyte Chemo-attractant Protein 1 (MCP-1/CCL2) supplementation was evaluated in this culture format and employed in dynamic co-culture conditions.

### Dynamic 3D co-culture

PHH spheroids were centrifuged at 100 xg, 1 min, and the supernatant was collected and centrifuged 1000 xg, 5 min to remove debris. PHH were resuspended in the dynamic 3D co-culture medium (50% of PHH-conditioned medium, 50% of fresh PHH medium without dexamethasone, supplemented with 50 ng/mL M-CSF and 50 ng/mL CCL2). Isolated monocytes were added to the resuspended PHHs in a monocyte-to-PHH ratio of 1:1 or 1:2. The cell mixture was inoculated in 30 mL spinners at a cell density of 1x10^5^ PHH/mL. After 4 days of culture, 25% of medium change was performed with fresh PHH medium without dexamethasone (final dexamethasone concentration: 37.5 nM, final FBS concentration: 5%). Isolated monocytes were incubated with CellTracker^TM^ Deep Red Dye (ThermoFisher Scientific) following manufacturer’s instructions before co-culture with PHH for both phagocytic activity and rAAV8-mCherry transduction evaluation.

### Flow Cytometry analysis

Samples from day 6 of 3D co-culture were collected, washed with PBS and incubated with TrypLE Express Enzyme (Thermo Fisher Scientific) for a maximum of 15 min. The resulting single cell suspension was washed three times in FACS buffer (PBS (-/-), 2% FBS) and incubated with the primary antibodies, for 45 min, at 4°C. Samples were washed and incubated with the viability dyes DAPI 1:1000 (Thermo Fisher, D3571) or 7AAD 1:50 (Biolegend, 420404), for 10 min, at room temperature. Samples were acquired in a BD FACSCelesta (BD Biosciences) and analyzed in the FlowJo (version 10.2, FlowJo) software package. Antibody references and dilutions are reported in Table 1.

**Table 1.** Antibodies used for flow cytometry and immunofluorescence.

|  | Brand | Catalog reference | Technique | Dilution |
| --- | --- | --- | --- | --- |
| CD45 | BD biosciences | 566041 | Flow Cytometry | 1:20 |
| CD45 | Biolegend | 304019 | Flow Cytometry | 1:20 |
| CD45 | BioLegend | 368512 | Immunofluorescence<br>Flow cytometry | 1:50<br>1:20 |
| MARCO | Invitrogen | 25-5447-42 | Flow Cytometry | 1:20 |
| MARCO | Sigma | HPA063793 | Immunofluorescence | 1:100 |
| VSIG4 | Invitrogen | 17-5757-42 | Flow Cytometry | 1:20 |
| CD16 | BD biosciences | 555406 | Flow Cytometry | 1:5 |
| CD163 | BioLegend | 326506 | Flow Cytometry | 1:20 |
| CD163 | Thermo Fisher | MA5-17716 | Immunofluorescence | 1:100 |
| CD206 | BioLegend | 321114 | Flow Cytometry | 1:20 |
| CD80 | BioLegend | 305220 | Flow Cytometry | 1:20 |
| HNF4α | abcam | ab41898 | Immunofluorescence | 1:100 |
| CK-18 | Sigma | F4772 | Immunofluorescence | 1:100 |
| Albumin | Santa Cruz | sc271605 | Immunofluorescence | 1:1000 |

CD45: Protein tyrosine phosphatase receptor type C (PTPRC), MARCO: Macrophage receptor with collagenous structure, VSIG4: V-set and immunoglobulin domain containing 4, CD16: Fc gamma receptor III, CD163: Cluster of Differentiation 163, CD206: Mannose receptor C-type 1, CD80: Cluster of differentiation 80, HNF4α : Hepatocyte nuclear factor 4 alpha, CK-18: Cytokeratin-18.

### 3D culture immunofluorescence characterization

PHH and PHH:KLC spheroids were fixed in 4% (w/v) paraformaldehyde with 4% (w/v) sucrose in PBS, for 20 min, RT. Samples were permeabilized and blocked with 0.1 % (v/v) Triton X-100 (TX-100) and 0.2 % (w/v) fish-skin gelatine (FSG) solution in PBS, for 30 min. The incubation of primary antibodies was performed for 2h at RT, followed by an incubation overnight at 4°C. After three washes with PBS, secondary antibody incubation was performed for 1h at RT. F-actin detection was performed by incubating samples with A488-conjugated Phalloidin (A12379, Thermo Fisher Scientific) 1:100 in PBS, for 20 min, after antibody incubation. Nuclei were stained with DAPI diluted in PBS (1:1000, Life Technologies), for 20 min. The samples were mounted onto coverslips in ProLong^TM^ Gold Antifade Mountant (Life Technologies). Antibodies and dye incubations were performed in a solution containing 0.125 % (w/v) FSG and 0.2% (v/v) Triton X-100. Images were acquired in a Zeiss 880-point scanning confocal microscope with Zeiss Zen 2.3 (black edition) or Leica Mica microhub microscope (confocal mode). Images were processed using ImageJ software. Antibody references and dilutions are reported in Table 1.

### Viral vector production and purification

An rAAV8 vector containing a GFP or mCherry reporter driven by the CMV promoter (rAAV8-GFP and rAAV8mCherry, respectively) was produced in suspension-adapted human embryonic kidney 293T cells (ATCC CRL-3216).[64] FreeStyle F17 expression medium, supplemented with 4 mM GlutaMAX (all Gibco), was used as a cell culture medium. At 2.0 x 10⁶ cell/mL, 293T cells were triple transfected using pHelper (Agilent), pRC8 (Addgene #112864, kindly provided by Dr. James M.Wilson, University of Pennsylvania) and in-house developed transgene using PEIpro (Polyplus) at 1:1.5 (w/w) DNA:PEI ratio. At 72h post transfection (hpt), the viruses were collected [65] and the rAAV8-mCherry particles were purified by affinity chromatography. Concentration and buffer exchange were performed against PBS containing CaCl_2_ and MgCl_2_ (14040091 Thermo Fisher Scientific). Samples were treated with DNase to eliminate free DNA contamination. Then, viral DNA was purified using the High Pure Viral Nucleic Acid Kit (Roche Diagnostics GmbH). Viral genomes (VGs) were quantified by quantitative PCR (qPCR) targeting CMV using the following prime-probe set: forward primer (ATATAGACCTCCCACCGTACAC), reverse primer (TGACGTCAATGGGAGTTTGTT) and probe-FAM (CCTACCGCCCATTTGCGTCAATG), at 0.5 µM and 0.25 µM final concentration, respectively. Thermal cycling was performed in a LightCycler 480 Instrument (Roche), at 95 °C for 10 min, followed by 40 cycles of 95°C for 10 sec, annealing at 62 °C for 30 sec, and extension at 72 °C for 6 sec (single acquisition), with a final cooling step at 40 °C. A linearized form of pAAV transgene was used as standard curve.

### Prototypical inflammatory stimuli on 3D cultures

After viability assessment and cell concentration determination, day 13 3D PHH and PHH:KLC cell cultures were collected and centrifuged at 100 xg, for 1 min, to sediment cell spheroids. The supernatant was collected and centrifuged at 400 xg, for 10 min to recover eventual myeloid cells not integrated in the PHH:KLC spheroids and pooled together with the PHH:KLC pellet. The supernatant was further centrifuged at 1000 xg, for 5 min, to remove cell debris. PHH and PHH:KLC pellets were resuspended in a medium composed of 50% of conditioned medium, from PHH or PHH:KLC spheroid cultures respectively, and 50% fresh PHH medium without dexamethasone (final dexamethasone concentration: 18.75 nM). Cells were plated in flat ultralow attachment (ULA) 24-well plates (5 x 10^5^ cell/mL, 125000 cell/well) and stimulated with 1 µg/mL Pam3CSK4, 1 µg/mL LPS or 10 µM ADU-S100 (final volume 500 μL/well). After 6h, samples were collected for RNA extraction. For ELISA evaluation, cells were stimulated with 0.1 µg/mL Pam3CSK4 or 0.1 µg/mL LPS for 6h, and the supernatant was collected, centrifuged at 10000 xg for 5 min (4°C) and stored at -80°C until further analysis.

### 3D culture transduction with rAAV

3D cultures were plated in 24-well ULA plates at the same cell density and medium composition used for incubation with prototypical stimuli. Cells were transduced with rAVV8-mCherry employing a multiplicity of infection (MOI) of 1x10^6^ VG/cell and samples collected at 6h, 24h, and 7 days. Samples from 6 and 24h were recovered for RNA extraction. Supernatants from 6 and 24h were recovered for ELISA analysis. Sample transduced for 7 days were collected and stored for immunofluorescence analysis (see below).

### Gene expression analysis

Total RNA was extracted with RNeasy Mini Kit (Qiagen), according to the manufacturer’s instructions. RNA was quantified in a NanoDrop 2000c (Thermo Fisher Scientific). Reverse transcription was performed using Sensiscript RT Kit (Qiagen), with a mix between anchored-oligo (dT)18 primer (10 µM) and random hexamers (Roche, 600 µM), or SuperScript^TM^ VILO^TM^ cDNA Synthesys Kit (Thermo Fisher Scientific).

Samples were mixed with molecular biology-grade water (Gibco), primers and LightCycler 480 SYBR Green I Master Kit (Roche). RT-qPCR reactions were performed in a LightCycler 480 Instrument II 384-well block (Roche) and quantification cycle values (Cts) and melting curves were determined using the LightCycler 480 Software version 1.5 (Roche). All data were analyzed relatively to the housekeeping gene *36B4* and presented as mRNA abundance or, by employing the 2^−ΔΔCT^ method for relative gene expression analysis,[66] as fold-change. Primer sequences are reported in Table 2.

**Table 2.**
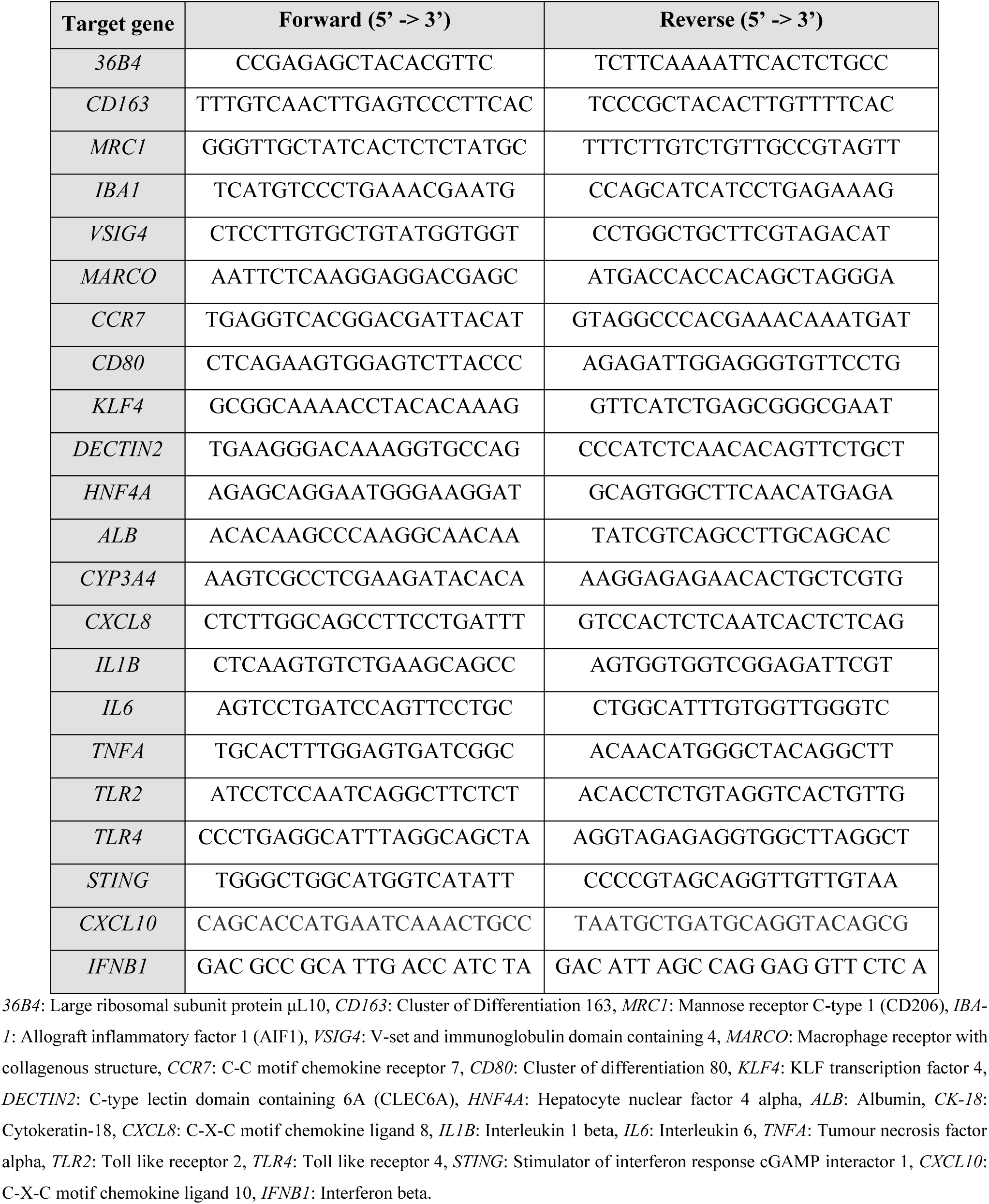
Primer sequences used for RT-qPCR.

### rAAV8-mCherry transgene expression analysis

The cDNA of transduced 3D cultures was quantified by droplet digital PCR (ddPCR), targeting mCherry. 3 µL of sample were added to a multiplex reaction containing target primer-probe set for mCherry (forward primer, AGTTCATGTACGGCTCCAA; reverse primer, GTCCTCGAAGTTCATCACG and, probe-FAM TCCCCGACTACTTGAAGCTGTC), and housekeeping gene TOP1 (forward primer, CTGTAGCCCTGTACTTCATCG; reverse primer, CTACCACATATTCCTGACCATCC and probe-HEX, CCTTCCTCCTTTTCATTGCCTGCTCT), at 0.90 µM for each primer and 0.25 µM for probe (final volume 22 µL). Droplets were generated by an Automated Droplet Generator (BioRad) and PCR reactions were performed in a C1000 Touch Thermal Cycler (BioRad), at 95°C for 10 min, followed by 40 cycles of 94°C for 30 sec, annealing/extension from 54 °C to 68 °C for 60 sec, and enzyme deactivation at 98°C for 600 sec, with a final cooling step at 4 °C. A QX200 droplet reader (Bio-Rad) using the QuantaSoft software package (Bio-Rad) was used to detect fluorescent signals in each droplet and calculate the concentration (copies/µL). mCherry concentration was normalized to TOP1.

### Enzyme-linked immunosorbent assay (ELISA)

The concentration of the inflammatory mediator IL-6 in the supernatant of 3D cultures stimulated with 1 µg/mL LPS, 1 µg/mL Pam3CSK4 and rAAV (MOI 1x10^6^ VG/cell) were measured by using the Human IL-6 ELISA Kit (R&D Systems) following manufacturer’s instructions.

### Phagocytic assay

Monocytes were plated in a 96-well plate and differentiated into M0-like macrophages as previously described. 3D cultures were plated in 24-well ULA plates as previously described. Cells were incubated with pHrodo™ Green *E. coli* BioParticles™ Conjugate for Phagocytosis (Thermo Fisher Scientific) at 250 μg/mL for 6h. To evaluate phagocytosis inhibition, a 3h pre-incubation with 10 μM Cytochalasin D (Focus Biomolecules) was performed. Samples were collected, washed three times with PBS and fixed in 4% (w/v) paraformaldehyde with 4% (w/v) sucrose in PBS, for 20 min, RT. Samples were stained with DAPI diluted in PBS (1:1000, Life Technologies), for 20 min. Images were acquired in Leica Mica microhub microscope and processed using ImageJ software.

### Statistical analysis

Statistical comparisons between two experimental conditions were performed using paired (when comparing the same PHH:KLC donor combination) or unpaired *t*-test (when comparing different donors), while for three or more experimental conditions one- and two-way ANOVA test, followed by Tukey’s multiple comparison test were used. Statistical significance was considered for *p* < 0.05, with \**p* < 0.05; \*\**p* < 0.01, \*\*\**p* < 0.001. GraphPad Prism v10.1.2 (GraphPad Software, San Diego, CA) was used for data analyses and graphics.

## Data availability statement

The original contributions presented in the study are included in the article/Supplemental Material, further inquiries can be directed to the corresponding author.

## Acknowledgments

The author(s) declare financial support was received for the research, authorship, and/or publication of this article. This work was funded by Boehringer Ingelheim International. We also acknowledge funding from FCT (MCTES (PT) through iNOVA4Health (UIDB/04462/2020 and UIDP/04462/2020); LS4Future (LA/P/0087/2020); 2022.12962.BD fellowship to I.R.G.( https://doi.org/10.54499/2022.12962.BD) and UI/BD/154874/2023 to I.P.C (https://doi.org/10.54499/UI/BD/154874/2023). The rAAV8-GFP and rAAV8-mCherry lots were produced and characterized by the Bioproduction Unit of iBET (www.ibet.pt), Portugal. Schemes in figures and graphical abstract were created with <u>BioRender.com</u>.

## Author Contributions

IRG: Conceptualization, Data curation, Formal Analysis, Investigation, Methodology, Visualization, Writing– original draft, Writing–review and editing. FA: Conceptualization, Data curation, Formal Analysis, Investigation, Methodology, Writing–review and editing. IPC: Data curation, Formal Analysis, Investigation, Methodology, Writing–review and editing. GD: Conceptualization, Methodology, Investigation, Writing– review and editing. SF: Data curation, Formal Analysis, Investigation, Methodology, Writing–review and editing. GS: Data curation, Formal Analysis, Methodology, Writing–review and editing. IS: Investigation, Writing – review and editing; ND: Data curation, Formal Analysis, Writing–review and editing. CF: Data Curation, Visualization, Formal Analysis, Writing–review and editing. PMA: Conceptualization, Funding acquisition, Resources, Writing–review and editing, Resources. UM: Conceptualization, Funding acquisition, Writing–review and editing, Resources. ASC: Conceptualization, Funding acquisition, Methodology, Supervision, Writing–review and editing. FDP: Conceptualization, Funding acquisition, Project administration, Writing–review and editing, Resources. CB: Conceptualization, Data curation, Funding acquisition, Methodology, Project administration, Supervision, Visualization, Writing–review and editing.

