## Supplementary information for "3D co-cultures of primary human hepatocytes and Kupffer-like cells to address innate immune responses to rAAV"

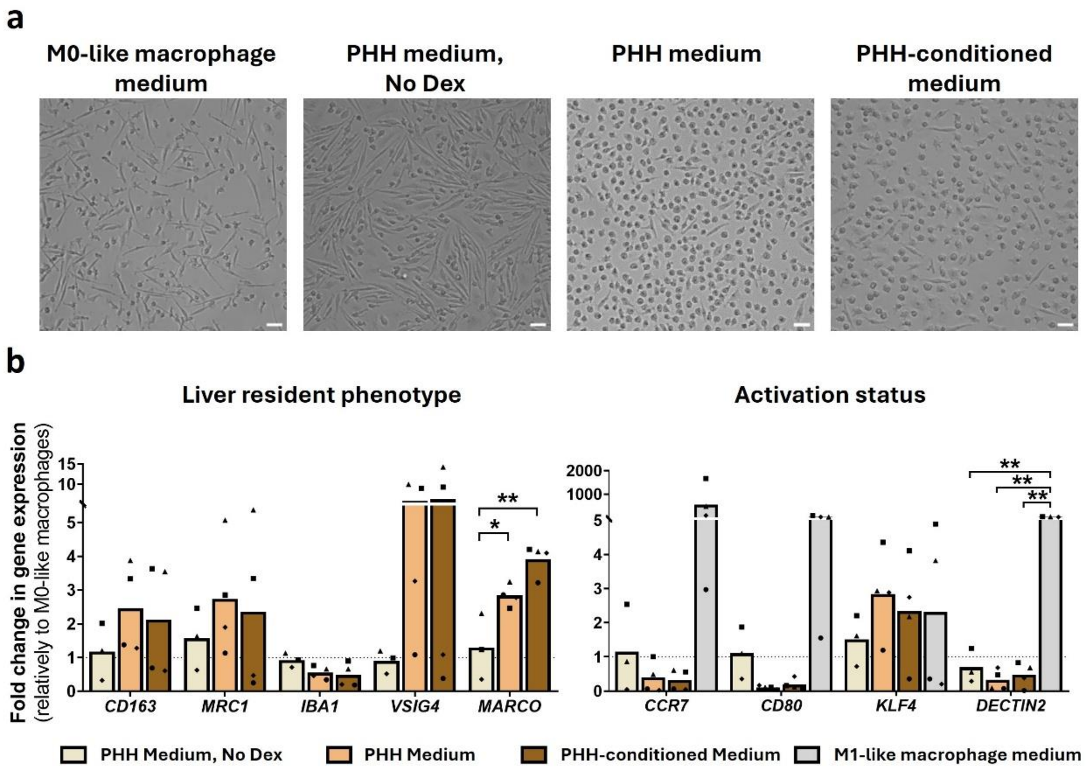


**Supplementary Figure S1 – Media composition evaluation for the differentiation of a Kupffer-like phenotype from 2D monocytes**

A) Morphology and B) gene expression evaluation of differentiated PBMC-derived monocytes in different media compositions for 6 days; scale bars: 50 µm. In B) Kupffer cell markers (left graph) and activation status (right graph) results are normalized for a housekeeping gene expression (36B4) and represented as fold change in respect to M0-like macrophages (non-stimulated macrophages, resting state). M1-like macrophages were employed as control of the myeloid population pro-inflammatory status (right graph). The symbols represent different PBMC donors; N ≥ 3. One-way ANOVA was employed for statistical analysis (Tukey's multiple comparisons test).

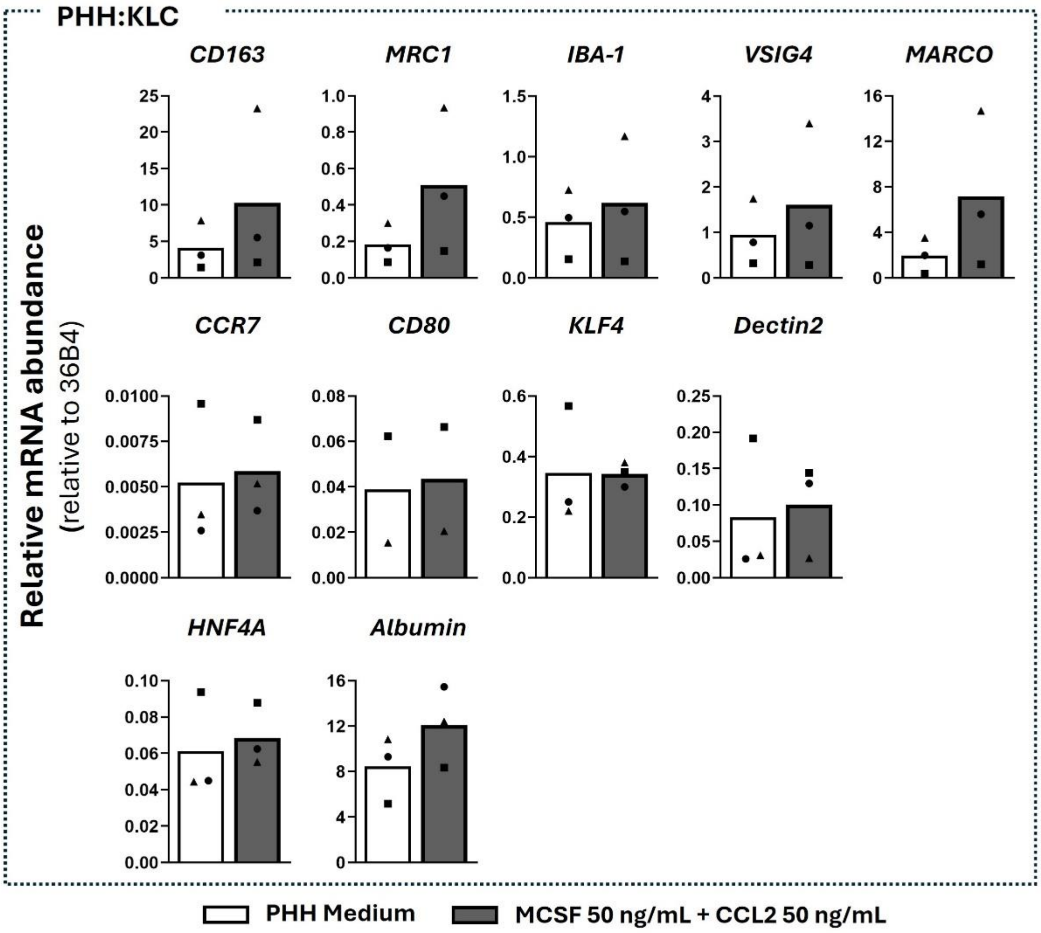


**Supplementary Figure S2 – Supplement addition effect on KLC and PHH populations in the 3D co-cultures**

A) Gene expression analysis of markers for Kupffer cell identity (upper graphs), activation status (central graphs) and hepatic identity (lower graphs) of day 6 static 3D co-cultures with or without M-CSF 50 ng/mL and CCL2 50 ng/mL supplementation. Data are expressed as relative mRNA abundance relative to 36B4. Symbols correspond to different PHH:KLC donor combinations. Paired T-test was performed for statistical analysis. N = 3 (CD80 N=2).

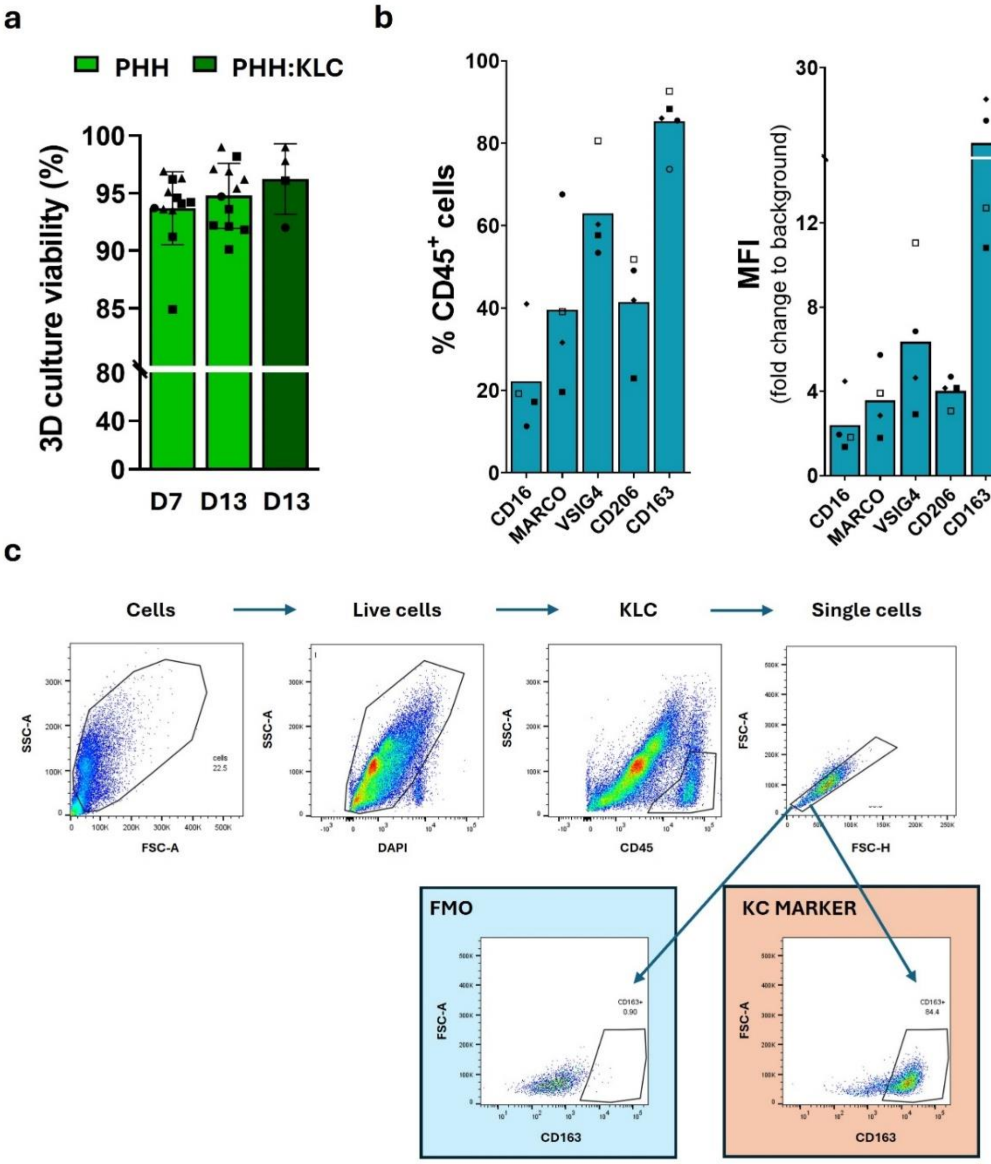


**Supplementary Figure S3 – Effect of KLC differentiation in 3D cultures and flow cytometry gating strategy**

A) Quantification of 3D cultures cell viability at day 7 and 13 for PHH monocultures (light green, day 7 and 13) and PHH:KLC co-cultures (dark green, day 13). Results are represented as mean and S.D of at least 3 independent experiments. Each symbol represents one biological replicate; different symbols represent different PHH donors that were employed in combination with different PBMC donors. One-way ANOVA was employed for statistical analysis. B) Flow cytometry evaluation of Kupffer-specific markers in PHH:KLC spheroids at day 13 of 3D culture. Results are expressed as percentage of positive cells (upper graph) and median fluorescence intensity (MFI, lower graph) detected inside of CD45+ population. Symbols correspond to different PHH:KLC donor combinations. N=4. Unpaired T-test was employed for statistical analysis. C) Gating strategy for the myeloid population identification in 3D co-cultures. After discarding debris, a DAPI+ sample was compared with a DAPI- to define the appropriate gate for live cells. Similarly, a CD45- and CD45+ were compared to define the gate for the KLC population. After selecting of the single cells in the myeloid cell population, we employed the Fluorescence Minus One (FMO) control to define the gate for each evaluated marker. The same gates were applied to all the performed co-cultures.

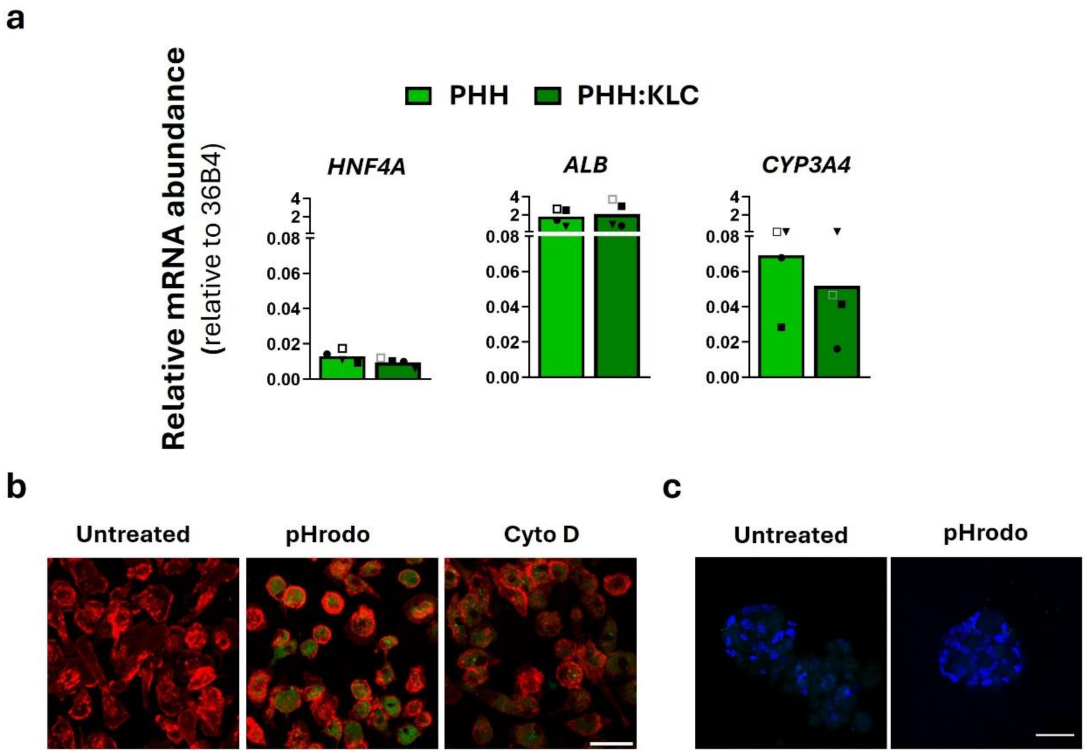


**Supplementary Figure S4 – Effect of KLC population on hepatic markers and phagocytic activity**

A) Evaluation of hepatic marker gene expression in PHH and PHH:KLC after 13 days of 3D culture. Data are expressed as relative mRNA abundance normalized to 36B4. N=4. Paired T-test was employed for statistical analysis. Different symbols correspond to different PHH:KLC combinations. B) Phagocytic activity assessment in 2D M0-like monocultures. Samples were incubated with pHrodo™ green BioParticles™ (250 µg/mL) for 6h with or without pre-incubation with 10 µM Cytochalasin D (Cyto D, phagocytosis inhibitor) for 3h. Cells were stained with CD45 (red). Scale bar = 50 µm. C) Phagocytic activity assessment in 3D PHH monoculture after 13 days of 3D culture. Samples were incubated with pHrodo™ green BioParticles™ (250 µg/mL) for 6h. Nuclei were stained with DAPI. Scale bar = 50 µm.

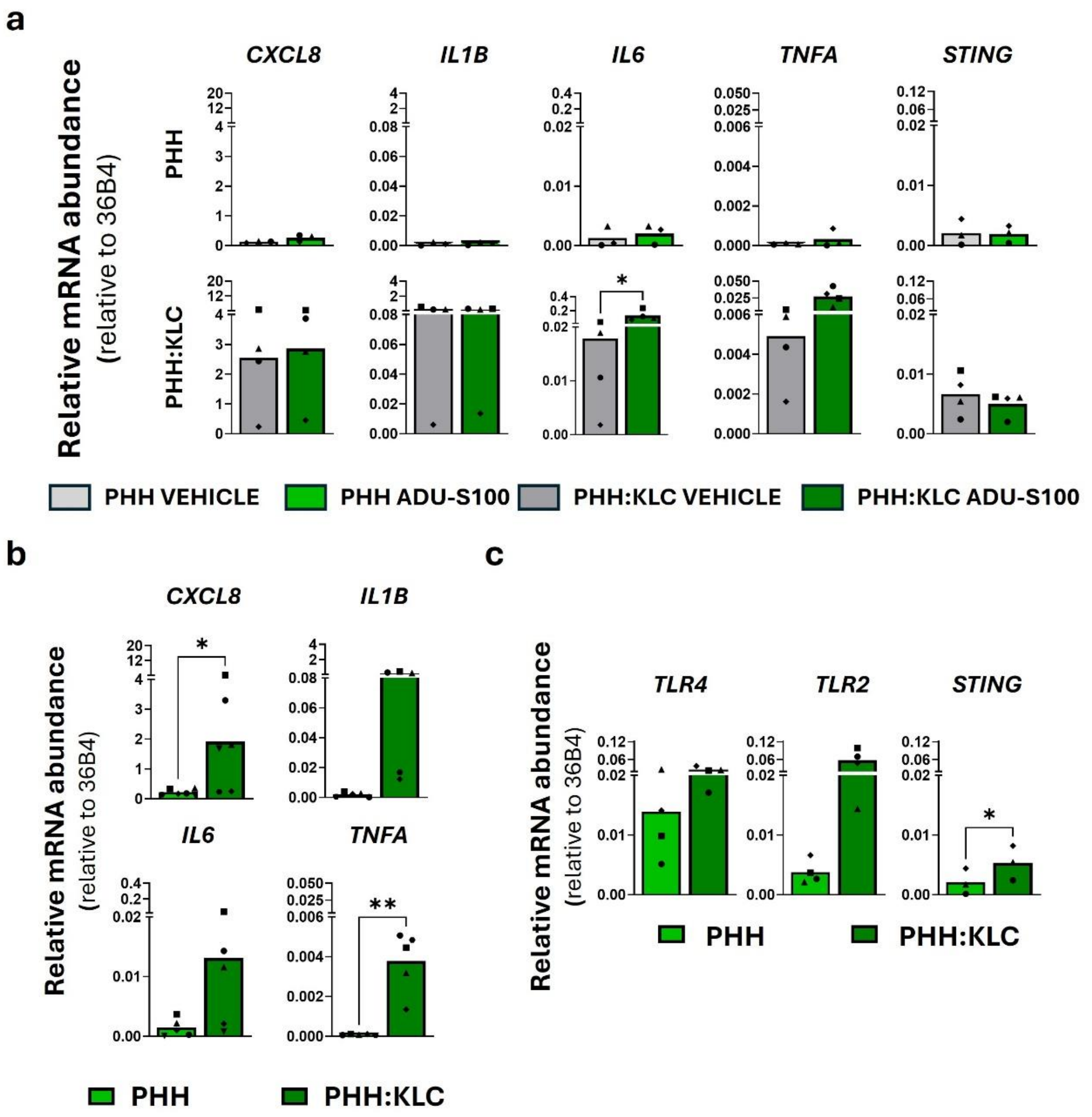


**Supplementary Figure S5 – 3D PHH, 3D PHH:KLC and M0-like response to stimuli and basal level comparison**

A) Gene expression evaluation of cytokines and STING after stimulation with ADU-S100 10 µM for 6h in PHH and PHH:KLC (13 days of 3D culture). Data are expressed as relative mRNA abundance relative to 36B4; N ≥ 3. Paired T-test was performed for statistical analysis. B) Cytokine and C) PRR gene expression evaluation in unstimulated PHH and PHH:KLC after 13 days of 3D culture. Data are expressed as relative mRNA abundance relative to 36B4; N ≥ 3. Paired T-test was employed for statistical analysis. In all graphs, symbols represent different PHH:KLC donor combinations.

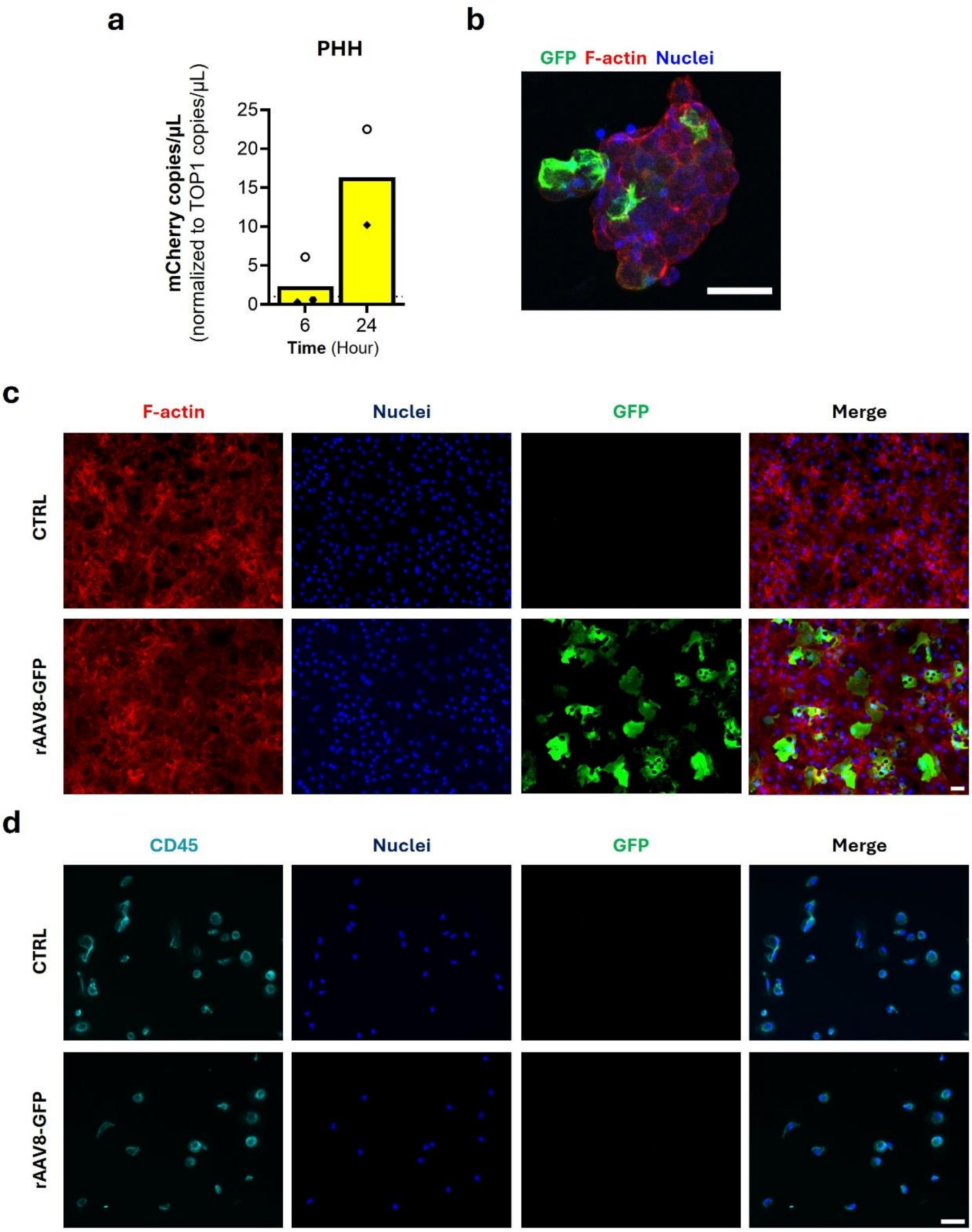


**Supplementary Figure S6 – PHH and macrophage susceptibility to rAAV8**

A) Evaluation of mCherry gene expression after PHH 3D culture transduction with rAAV8-mCherry for 6 and 24h. Results are expressed as fold change in respect to the vehicle condition. N = 3 for 6h, N = 2 for 24h. B) Immunofluorescence confocal microscopy evaluation of PHH 3D co-cultures transduced with rAAV8-GFP (MOI $1x10^6$ VG/cell). Scale bar: 50 µm. C) and D) Immunofluorescence microscopy of C) 2D PHH and D) M0-like macrophages to evaluate AAV8 transduction. Cells were transduced for 7 days with rAAV8-GFP (MOI $1x10^6$ VG/cell). In A) PHH were stained for F-actin (phalloidin, red) and GFP (green). In D) M0-like macrophages were stained for CD45 (light blue) and GFP (green). Nuclei were stained with DAPI (blue). Scale bars: 50 µm.
